# Functional study of two flexible regions of the hepatitis E virus ORF1 replicase

**DOI:** 10.1101/2025.10.22.683852

**Authors:** Léa Mézière, Sonia Fieulaine, Claire Montpellier, Martin Ferrié, Thibault Tubiana, Gabriel Vanegas Arias, Stéphane Bressanelli, Laurence Cocquerel, Cécile-Marie Aliouat-Denis

**Affiliations:** Univ. Lille, CNRS, INSERM, CHU Lille, Institut Pasteur de Lille, U1019-UMR 9017-CIIL-Center for Infection and Immunity of Lille, F-59000 Lille, France; Université Paris-Saclay, CEA, CNRS, Institute for Integrative Biology of the Cell (I2BC), Gif-sur-Yvette, France

**Keywords:** HEV, replication, AlphaFold2, linker, genotype 3, replicon, luciferase, mutagenesis, cleavage, flexible hinge

## Abstract

Hepatitis E virus (HEV), like other positive-sense RNA viruses, encodes a multidomain protein essential for replication, termed ORF1. However, the number, organization, and functions of its domains remain debated. Using AlphaFold2-based structural modeling, we investigated two structurally disordered regions with potential regulatory functions: (i) a 16-residue linker between the Helicase (Hel) and RNA-dependent RNA polymerase (RdRp) domains, proposed as a cleavage site and/or a flexible hinge, and (ii) the RdRp C-terminal tail, suggested to modulate polymerase activity through conformational plasticity.

We performed mutagenesis of the Hel/RdRp linker and analyzed the impact of mutations on ORF1 maturation, subcellular localization, and replication efficiency. Using an extensive antibody panel combined with precise protein sizing, we found that ORF1 is predominantly expressed as a full-length protein with an apparent molecular weight of ∼235 kDa by SDS-PAGE, together with several low-abundance truncated products, including the previously described HEV-derived SMAD activator (HDSA) fragment. These results suggest that ORF1 likely undergoes post-translational modifications and partial maturation by cellular proteases and/or spontaneous truncations in exposed regions. Importantly, Hel/RdRp linker mutations did not alter the ORF1 expression profile or subcellular localization, arguing against cleavage within this region. However, substitutions of conserved residues in the linker strongly impaired HEV replication, highlighting the functional importance of this disordered segment for viral genome replication.

Similarly, deletions or substitutions within the last 20 C-terminal RdRp residues abolished or severely impaired HEV replication. This demonstrates that the conformational flexibility of the RdRp C-terminal segment is likely critical for ORF1 function, e.g. for polymerase activity.

In conclusion, although ORF1 likely undergoes tightly regulated processing, cleavage is unlikely to occur within the Hel/RdRp linker. Nevertheless, this segment and the conformational dynamics of the RdRp C-terminus emerge as key regulatory elements required for efficient HEV replication, pointing to novel mechanistic layers of control in the HEV replication process.

## Introduction

Hepatitis E virus (HEV) is the leading cause of acute hepatitis in humans. The WHO estimates 20 million HEV infections worldwide leading to 3.3 million symptomatic cases and 70,000 deaths (WHO, 2023; [1]). A lack of standardized reporting makes it difficult to measure the current burden of hepatitis E which is likely under-estimated [2]. HEV causes a significant global health burden, not only in low-to-middle income countries following a fecal-oral mode of contamination, but also in high-income countries as a food-borne pathogen to be surveilled in priority [3,4]. HEV has an important zoonotic potential, it was recently ranked 6th among the top 10 zoonotic viruses presenting the greatest risk of spillover and spread to humans [5], and was very recently included on the list of Public Health Emergencies of International Concern (WHO, Health Emergencies Program, June 2024).

Hepatitis E is generally self-limiting in 3 to 8 weeks with a case fatality rate of 0.5-3% in young adults. However, fulminant hepatitis may occur in pregnant women and result in up to 30% mortality [6]. Hepatitis E can also become chronic in immunosuppressed patients and may lead to fibrosis, cirrhosis and extra-hepatic manifestations such as neurological, pancreatic or kidney disorders. Despite its important impact on human health, no specific HEV treatment is available. An anti-HEV vaccine has been licensed in China and more recently in Pakistan [7].

The four genotypes that are responsible for the majority of hepatitis E human cases belong to the *Paslahepevirus* genus [8]. Genotypes 1 and 2 only infect human beings through fecal-oral route, generally *via* contaminated water and may cause extensive acute hepatitis E epidemics in low-income countries. Genotypes 3 and 4, responsible for chronic hepatitis in immunocompromised patients, spread according to a sporadic transmission mode, in industrialized countries and are transmitted zoonotically through the consumption of raw or undercooked pork or game meat. Recently, viruses isolated from rats and belonging to the *Rocahepevirus* genus have been recognized to elicit infection in immunocompromised as well as immunocompetent individuals [9]. Vertical transmission from mother-to-child or to patients after blood transfusion have also been reported [10].

HEV is a small quasi-enveloped virus. Its 7.2kb positive single-stranded RNA genome encodes 3 proteins named ORF1, ORF2 and ORF3. Structural proteins are ORF2, that corresponds to the viral capsid and ORF3, a small phosphoprotein involved in viral particle secretion. The ORF1, also known as the HEV replicase, is a 185-194 kDa non-structural polyprotein consisting of several domains essential for genome replication. Recently, AlphaFold2 (AF) modeling corrected the former assignment of domains in ORF1 protein. First, it confirmed and delineated the limits of the C-terminal enzymes, domains X, helicase and polymerase [11,12], whose enzymatic activities have been confirmed by several studies (reviewed in [13,14]). Then, it further showed that the two previously assigned N-terminal methyltransferase and Y domains were better described as a single MetY domain. The existence of the MetY domain was supported by a high AF confidence score (predicted local distance difference test, pLDDT > 80) and a strong homology with its counterpart nsP1 in Alphaviruses [11,12]. Finally, it revealed that ORF1 does not contain any protease domain [11,12,15]. Instead, a domain that folds as a fatty acid binding domain is found downstream the MetY domain (residues 515-707) [11,12,16]. Nevertheless, the region initially supposed to be a papain-like cysteine protease (PCP) domain (residues 434–592) still raises controversies in terms of enzymatic activity and/or function [11,15]. Recently, it has been demonstrated that this region in ORF1 is necessary for viral replication [17], likely through a hexa-Cys motif (CxCx_11_CCx_8_CxC in region 457–483) embedded in the MetY domain, that is likely to bind divalent cations, most probably zinc [15,17,18]. Moreover, while it is now clear that there is no protease in HEV ORF1 [15], there is conflicting evidence as to whether the HEV ORF1 polyprotein requires proteolytic processing to facilitate the functions of its subdomains. Indeed, some studies suggest that ORF1 maturation by cellular proteases is necessary for subdomain functionality, while others argue that ORF1 functionality is independent of proteolytic cleavage (reviewed in [19]). Thus, whether the ORF1 functions as a full-length flexible protein or requires processing by one or more cellular proteases to fulfill its roles during the viral cycle remains an open question [19].

Here, we investigated the function of two structurally disordered regions (pLDDT < 50) that are interspersed between well-structured ORF1 domains and could play an important role in the subtle regulation of ORF1 enzymatic activities at different stages of the HEV lifecycle. Indeed, the flexibility of ORF1 polyprotein could be instrumental in its functional regulation. We first studied a 16 amino acid linker region located between the Hel and RdRp domains (**Fig 1a**, top), which could confer relative flexibility to these domains and constitute a cleavage site accessible to cellular protease(s). We also studied another disordered region of 19 amino acid residues located at the C-terminal tail of the RdRp domain (**Fig 1a**, bottom). The flexibility of this region and subsequent alternate folding in a region of the RdRp critical for RNA synthesis could lead to down-regulation of the polymerase and thus, to the fine control of its activity during the HEV lifecycle. Site-directed mutagenesis was conducted in both regions and the impact of mutations was analyzed by studying the ORF1 processing and subcellular localization, as well as the efficiency of viral replication.

**Fig 1.**
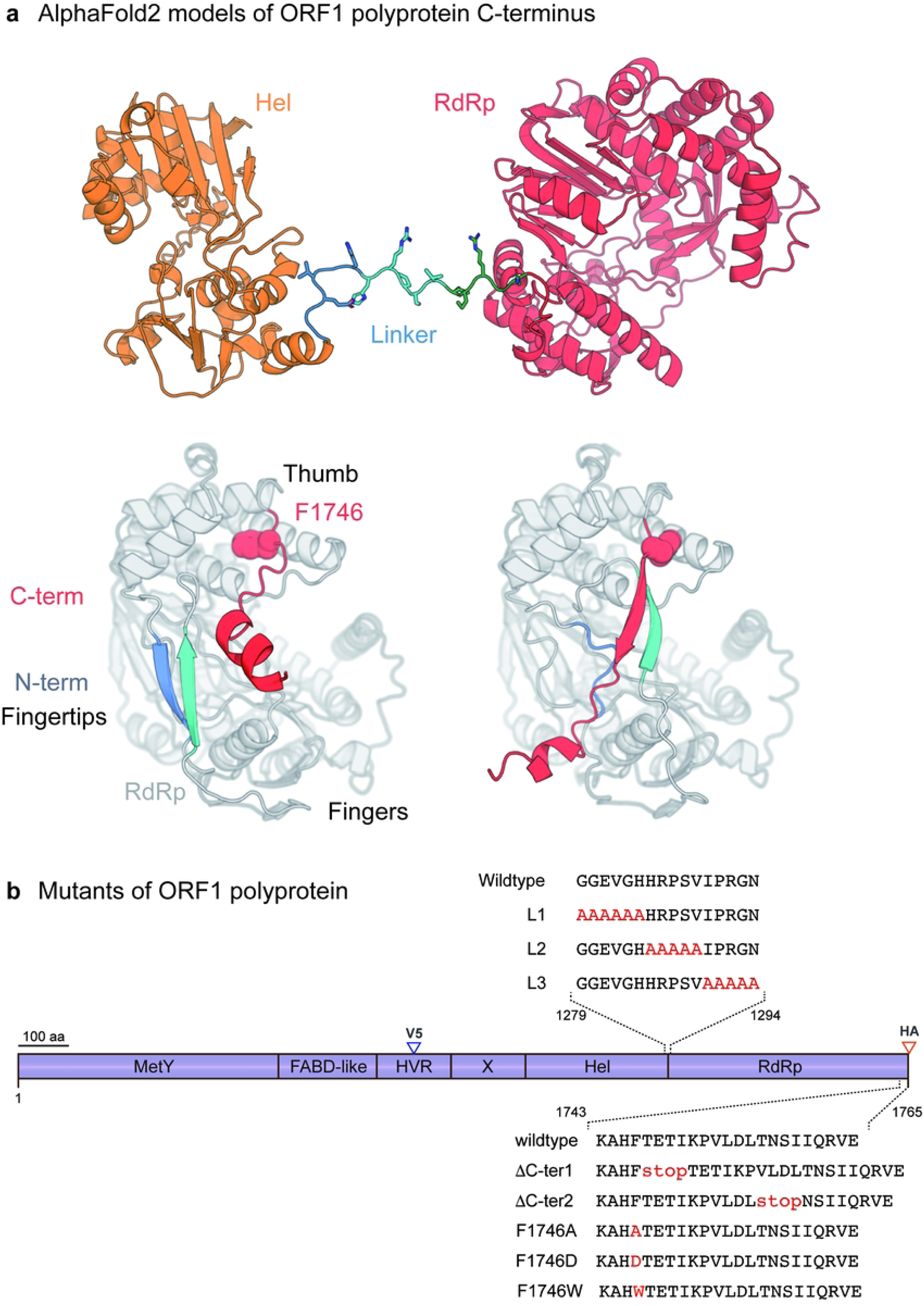
AlphaFold2 models of HEV ORF1 polyprotein C-terminus and localization scheme of mutations and epitope tag insertions. (a) AlphaFold2 models of HEV ORF1 polyprotein C-terminal domains. Top, C-terminal 764 residues (gt3 Kernow C-1 p6 HEV strain numbering). Orange, Hel domain; blue to cyan to green, linker; red, RdRp. Bottom, two alternate models of RdRp fold. The 20 C-terminal residues in alternate conformations are colored in red, with F1746 displayed as spheres. Two segments from the RdRp N-terminus also in alternate conformation are colored in blue and cyan; Left, these two segments pair in a beta-sheet, while the C-terminal segment is not included in the fingertips and F1746 is deeply anchored in the thumb. Right, F1746 is more loosely anchored in the thumb and the C-terminal segment is embedded in the fingertips between the two blue and cyan segments. (b) Schematic representation of the ORF1 polyprotein. ORF1 functional domains are designated as methyltransferase and Y domain (MetY), fatty-acid binding domain (FABD-like), hypervariable region (HVR), macro-or X-domain (X), helicase (Hel), RNA-dependent RNA polymerase (RdRp). Insertions of V5 and HA epitopes are positioned in the HVR and at the C-terminus of ORF1, respectively (blue and red arrowheads, respectively). Amino acid sequences of interest in this study are detailed in the Hel-RdRp linker region (top) and at the C-terminus (bottom) of RdRp to highlight the position of mutated amino acid residues (red) compared to the wildtype sequence.

## Material and methods

### Cell culture

The Huh-7-derived H7-T7-IZ cells stably expressing the T7 RNA polymerase [20], kindly provided by Pr. R. Bartenschlager (University of Heidelberg, Germany), were maintained in Dulbecco’s modified Eagle’s medium (DMEM; Thermo Fisher Scientific) supplemented with 10% foetal bovine serum (FBS), 1% non-essential amino acid (NEAA; Thermo Fisher scientific) and 50 μg/ml of Zeocin (InvivoGen). They were used for the transfection of the T7 promoter-driven pTM expression vectors. The PLC/PRF/5 (CRL-8024) derived PLC3 cells were characterized as highly replicative and productive cell line for HEV [21]. PLC3 cells were grown in DMEM supplemented with 10% FBS and 1% NEAA to study HEV replication efficacy.

### Plasmids and transfection

All experiments were performed with ORF1 from HEV gt 3 Kernow C-1 p6 strain (GenBank accession number JQ679013). The pTM-ORF1 plasmid, kindly provided by Dr. J. Gouttenoire (University of Lausanne, Switzerland) contains the full-length sequence of the ORF1 protein [22]. In a previous study, a V5 epitope has been inserted in the ORF1 hypervariable region (ORF1-V1, [23]). This construct served as a backbone to mutate the linker region (G_1279_-N_1294_) lying between the ORF1 helicase and RNA-dependent RNA polymerase domains (**Fig 1b**). To generate L1, L2 and L3 linker mutants, 3 blocks of amino acid residues (G_1279_-H_1284_, H_1285_-V_1289_, I_1290_-N_1294_, respectively) were mutated to alanine residues by using the fusion PCR technique between the NdeI/NsiI restriction sites and the indicated primers (**Table 1** and **Fig 1b**). To generate the V5-HA double-tagged-ORF1, an AflII/MluI restriction digest was performed to replace the C-terminal ORF1 region of L1-L3 mutants by the HA-tagged C-terminus of the pTM-ORF1-H1 described previously [23].

**Table 1.**
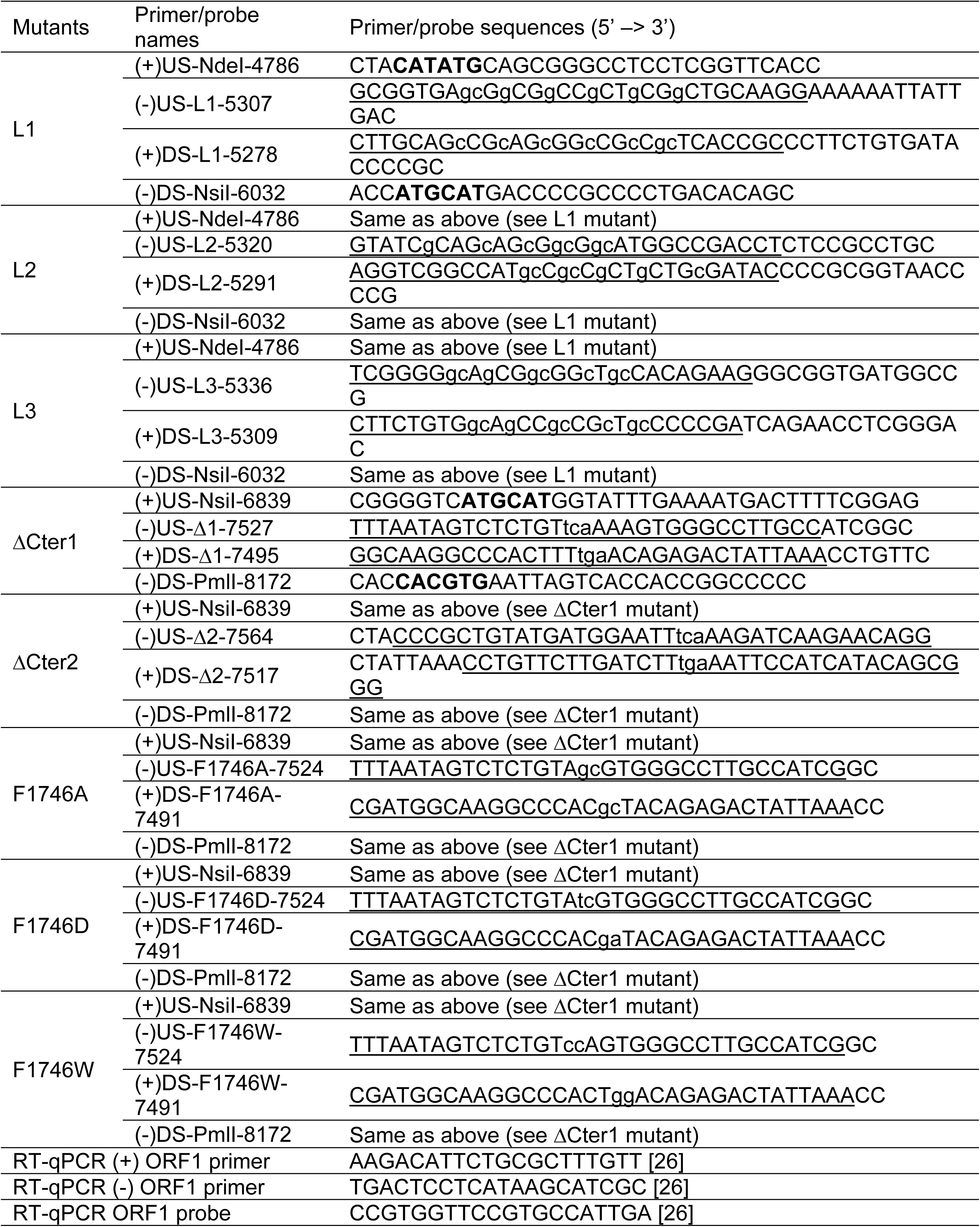
Primer/probe sequences used in the fusion PCR to construct the different ORF1 mutants as well as in RT-qPCR experiments to quantify replicon RNA. (+): forward primer, (-): reverse primer, US: upstream fragment, DS: downstream fragment, L1 = mutant 1 in the Hel-RdRp linker region, L2 = mutant 2 in the Hel-RdRp linker region, L3 = mutant 3 in the Hel-RdRp linker region. Restriction site sequences are highlighted in bold. Small letters symbolized inserted mutations. Overlapping regions used in the PCR fusion to anneal the upstream and downstream PCR products are underlined.

H7-T7-IZ cells were transfected with T7 promoter-driven pTM expression vectors. The pTM plasmids expressing V5-or V5- and HA-tagged wiltype (WT) or mutated (L1, L2, L3) ORF1 were transfected using the Viafect transfection reagent (Promega). After 16-24h of incubation, transfected cells were lysed or fixed and analyzed by Western blotting (WB) / immunoprecipitation (IP) or immunofluorescence (IF), respectively.

The plasmid pBlueScript SK (+) carrying the DNA of the full-length genome of HEV gt 3 Kernow C-1 p6 strain was kindly provided by Dr. S.U. Emerson. The HEV p6-wt-GLuc replicon, constructed from the HEV gt 3 Kernow C-1 p6 strain was also obtained from Dr. Emerson. This replicon possesses a Gaussia Luciferase (GLuc) reporter gene that substitutes the 5’ part of the ORF2 gene and most part of the ORF3 gene [24,25]. A p6-GAD-GLuc mutant replicon in which the ORF1 polymerase active site GDD was mutated to GAD to prevent any replication was used as a negative control [25]. Directional restriction digests (NdeI/NsiI) were performed to replace the wildtype Hel-RdRp linker sequence by the L1-L3 mutated sequences into the p6-GLuc replicon in order to assess the replication efficiencies of these linker mutants.

Mutations in the RdRp C-terminal region were introduced into the p6-Gluc replicon by using the fusion PCR technique between the NsiI/PmlI restriction sites and the indicated primers (**Table 1** and **Fig 1b**). Five ORF1 mutants were studied for their replication efficiencies: ΔCter1 for which a stop codon was introduced downstream of F1746, ΔCter2 for which T1757 was mutated to a stop codon, F1746A, F1746D and F1746W for which F1746 was mutated to A, D and W, respectively. Mutations and epitope insertions were verified by DNA sequencing for all constructs.

Capped genomic HEV RNAs were prepared with the mMessage mMachine kit (Life technologies) and delivered to PLC3 cells by electroporation using the Gene Pulser Xcell apparatus (Bio-Rad), as described in [21]. Electroporated PLC3 cells were seeded in 24-well plates and incubated at 37°C under 5% CO_2_.

### Western blotting

H7-T7-IZ cells were lysed during 20 min at 4°C with lysis buffer (10 mM of Tris-HCl (pH 7.0), 150 mM of NaCl, 2 mM of EDTA, 0.5% Triton X-100) supplemented with protease inhibitors (cOmplete^TM^ protease inhibitor cocktail, Roche) and 1 mM PMSF (Roche). Then, cells debris were eliminated by centrifugation at 6000 rpm during 10 min at 4°C. Cells lysates were stored at -20°C until analysis. Before loading, cytoplasmic extracts were denatured in Laemmli buffer/DTT 200 mM at 70°C for 10 min. Then, proteins were separated by 8% bis-acrylamide SDS-PAGE, and transferred to nitrocellulose membranes (Hybond-ECL, Amersham). The proteins of interest were detected by specific primary antibodies and corresponding horseradish peroxidase-conjugated secondary antibodies (**Table 2**). Membranes were washed six times with washing buffer (1X-PBS, 0.2% Tween20) between each step. Finally, nitrocellulose membranes were incubated with SuperSignal™ West Pico plus or Pierce ECL Western blotting substrate (Thermo Fisher scientific), and revealed by using a developer (Amersham) or ImageQuant™ 800 western blot imaging system (Cytiva). Page Ruler and ProSieve Quadcolor protein markers were purchased respectively from Thermo Fisher scientific and Lonza.

**Table 2.**
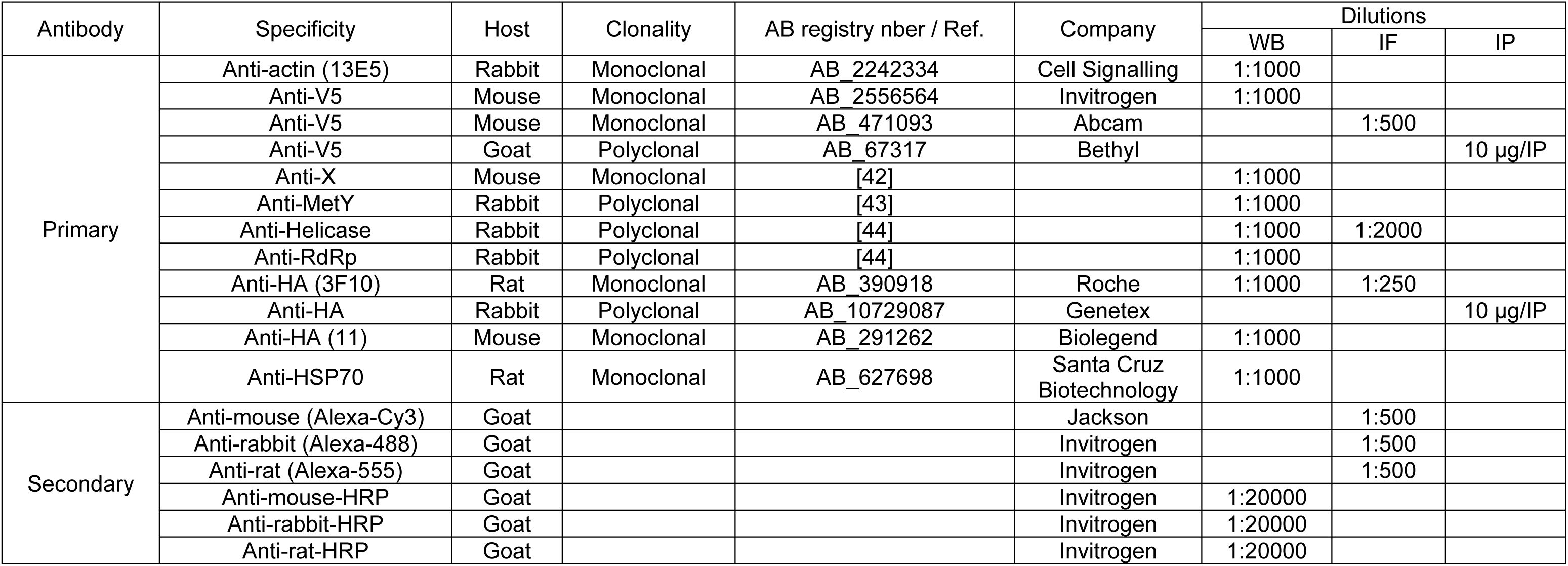
Primary and secondary antibodies used in Western blot (WB), immunofluorescence (IF) and immunoprecipitation (IP) experiments.

### Chemicals

MG132, a tripeptide aldehyde inhibitor of proteasome, and NVP-HSP-990, a selective inhibitor of heat shock protein 90 (HSP90), were purchased from MedChemExpress. They were dissolved in DMSO (Sigma).

### Indirect immunofluorescence

H7-T7-IZ cells transfected with wildtype or mutant ORF1 pTM constructs were seeded on glass coverslips in 24-well plate and fixed during 20 min at room temperature (RT) with 3% paraformaldehyde (PFA) at 24h post-transfection. Then, cells were washed thrice with 1X-PBS, followed by cell membrane permeabilization with 0.5% Triton X-100 diluted in 1X-PBS during 30 min at RT. To block nonspecific sites, cells were incubated with 10% goat serum in 1X-PBS for 30 min at RT. Then, target proteins were stained with specific antibodies for 30 min at RT, followed by three wash and an incubation with fluorochrome-conjugated secondary antibodies for 30 min at RT in the dark (**Table 2**). Nuclei were stained with DAPI (4’,6-diamidino-2-phenylindole, 1:500 in 10% goat serum/1X-PBS). The cells were washed thrice in 1X-PBS and coverslips were mounted with Mowiol (Calbiochem) on glass slides. The microscopic observation was conducted using LSM 880 confocal laser scanning microscope (Zeiss) (63x oil). Fiji software was used to analyze images.

### Immunoprecipitation (IP)

Antibodies directed against V5 (Bethyl) or HA tags (Genetex) were bound to magnetic beads using the Dynabeads antibody coupling kit (Thermo Fischer Scientific), overnight at 37°C, following the manufacturer’s instructions. Beads were then washed and incubated for 1h at RT with cytoplasmic extracts from transfected H7-T7-IZ cells. Next, beads were washed six times with 1X-PBS/0.5% NP40. Then, beads were suspended in Laemmli buffer/DTT 200 mM and heated at 70°C for 10 min. Eluted samples were analyzed by WB using anti-V5 (Invitrogen) or anti-HA (clone 3F10, Roche) antibodies, as described above.

### Luciferase assay

PLC3 cells were electroporated with p6-Gluc-ORF1-WT, p6-GLuc-ORF1-Mut-Cter, p6-Gluc-ORF1-Mut-link and p6-Gluc-ORF1-GAD replicons and incubated for 6 days at 37°C with 5% of CO_2_. Culture supernatants, containing the cell-secreted-luciferase, were sampled at 8, 24, 72, 96, 120 and 144 hours post-electroporation. Samples were stored at -20°C until analysis. The supernatants were diluted at 1:100 in 1X-Passive lysis buffer (Promega), then 5 μL were transferred into a 96-well plate before reading. The luminometer (Centro XS3 LB960, Berthold Technologies) was programmed to deliver 20 μL of luciferin substrate (Renilla Luciferase Assay System, Promega) per well, and to measure the light signal for 5 s. Luciferase activities measured at each time point were normalized by the activities measured at 8 h post-electroporation.

### Quantification of replicon RNA

After electroporation of PLC3 cells with the p6 luciferase replicons, total RNA was extracted from cells (NucleoSpin RNA plus kit, Macherey-Nagel) at 6 days post-electroporation. One-step qPCR assay was performed using 5 μL of RNA and Takyon Low rox one-step RT probe master mix (UFD-LPRT-C0101, Eurogentec) with specific ORF1 primers and probes designed against genomic RNA according to previously published literature (**Table 1**; [26]), and using a Quantstudio 3 (Applied Biosystems). In order to quantify the HEV genome, a standard curve was prepared by diluting the *in vitro*-transcribed HEV p6 plasmid in total RNA extracted from mock electroporated PLC3 cells.

### Statistical analyses

Statistical analyses were performed using GraphPad Prism 9.5.0 software. For comparing replication kinetics, 2way ANOVA and Dunnett’s multiple comparisons tests were performed. For comparing percentage of replication, the non-parametric Kruskal-Wallis test was used. Groups were compared to the WT control. A test was declared statistically significant for any p value below 0.05. Data are presented as mean ± SD. **p<0.01, ***p<0.001, ****p<0.0001.

### Molecular modeling

We used our previously generated AlphaFold2 models of pORF1 from HEV gt 3 Kernow C-1 p6 strain [11]. The models displayed in **Figs 1a** and **S1a Fig** were generated with PyMol.

## Results

### Characterization of the linker region between the Helicase and RdRp domains of HEV ORF1

#### Impact of mutations in the Hel/RdRp linker region on the expression profile and maturation of the ORF1 polyprotein

Structural modelling using AF software has recently identified a 16-amino acid linker region located between the Hel and RdRp domains [11,12]. This disordered region could confer relative flexibility between these 2 domains and/or constitute a cleavage site accessible to protease(s) (**Fig 1a**, top). To gain new insights into the mechanistic regulation of ORF1 activity, we first investigated the importance of this disordered region on HEV ORF1 maturation. The 16-amino acid linker region was mutated, i.e. 3 blocks of amino acid residues (G_1279_-H_1284_, H_1285_-V_1289_, I_1290_-N_1294_) were mutated to alanine residues in order to generate L1, L2 and L3 linker mutants, respectively (**Fig 1b**, top). These constructs were engineered in a backbone plasmid in which a V5 epitope was inserted in the ORF1 hypervariable region (**Fig 2a**, blue arrowhead), as previously described [23]. A heterologous expression system was first used to analyze the expression profiles of wildtype and mutant ORF1 proteins. To this end, T7 promoter-driven pTM vectors expressing wildtype or mutated ORF1 were transfected in the Huh-7-derived H7-T7-IZ cells stably expressing the T7 RNA polymerase, as previously described [23] (**Fig 2**).

**Fig 2.**
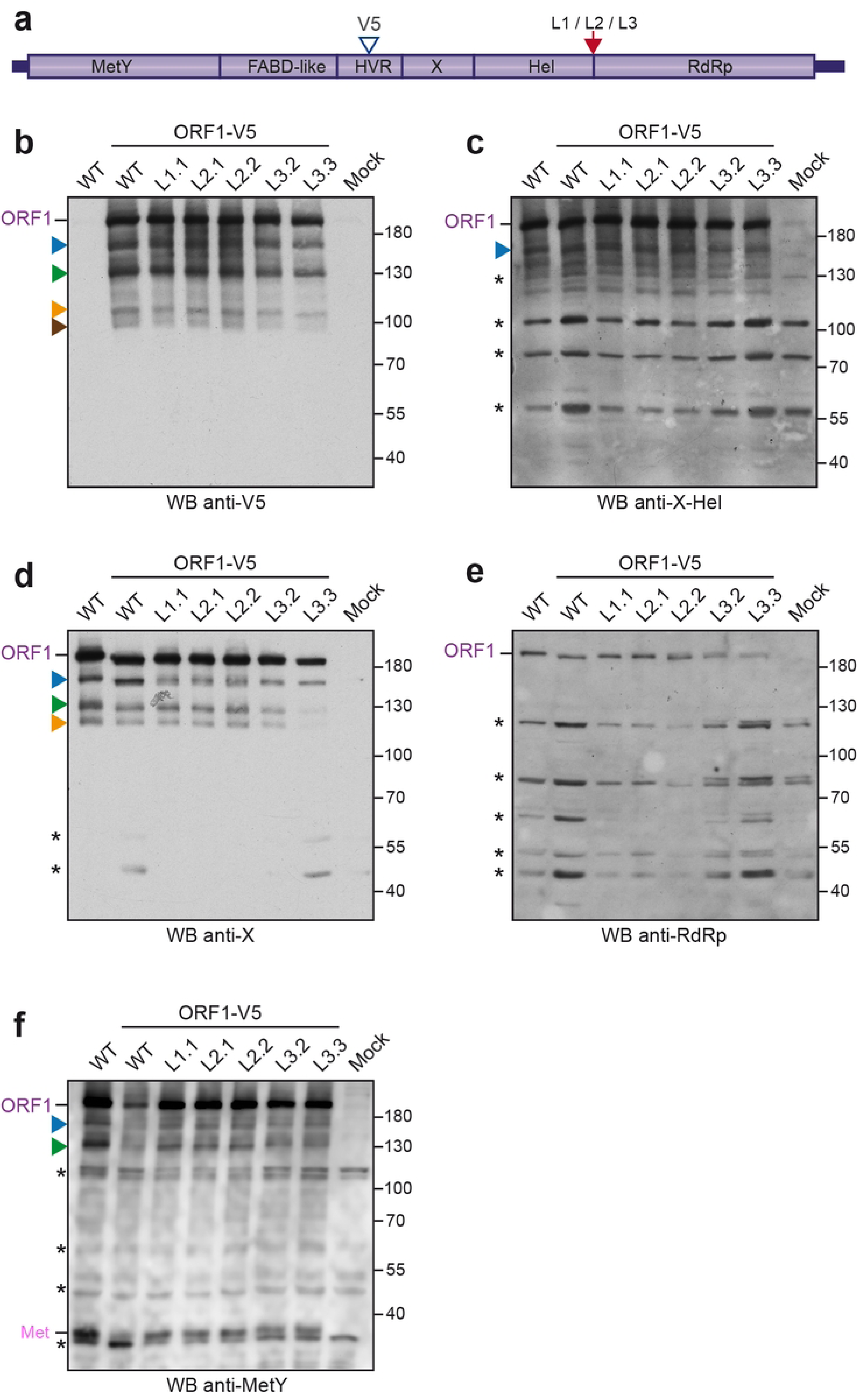
Western blot analysis of H7-T7-IZ cells expressing V5-tagged wildtype and mutant ORF1 proteins in the Hel/RdRp linker region. (a) H7-T7-IZ cells were transfected with a pTM plasmid expressing the untagged (WT) or V5-tagged wildtype ORF1 protein (V5 blue arrowhead) or the ORF1 harboring mutations in the Hel/RdRp linker region (L1, L2, L3, red arrow). Total cell lysates were collected in the presence of protease inhibitors at 24 h post-transfection and analyzed by WB. Mock-transfected cells served as a negative control (Mock). Immunoblots were probed either with antibodies directed against the V5 epitope (b), the X-Hel domain (c), the macrodomain X (d), the RdRp domain (e) or the MetY domain (f). The full-length ORF1 protein (ORF1) and the Met domain (Met) are indicated on the left. Proteins detected at lower molecular weights are also indicated (blue, green, orange and brown arrowheads). Molecular weight markers are indicated in kilodaltons on the right. Two clones have been analyzed per construct except for the untagged wildtype (WT) and L1 ORF1 mutant. Asterisks indicate non-specific bands.

At 24 h post-transfection, cell lysates were prepared and analyzed by Western blot (WB) using anti-V5 tag, anti-X-Hel domain, anti-X domain, anti-RdRp and anti-MetY domain antibodies (**Fig 2b-f**). As expected, no ORF1 protein expression was detected in non-transfected cells (Mock, last lanes). Moreover, the specificity of the anti-V5 antibody was confirmed by the absence of signal in cells expressing the wildtype untagged ORF1 protein (WT) (**Fig 2b**, first lane). The anti-V5 WB revealed several specific bands corresponding to the full-length ORF1 protein as well as proteins with smaller molecular weights (**Fig 2b**, ORF1, blue, green, orange and brown arrowheads). The recognition patterns remained the same irrespective of whether cells expressed V5-tagged wildtype and mutant ORF1 proteins. The full-length ORF1 protein was also detected by all tested antibodies recognizing X-Hel, X, RdRp and MetY domains (**Fig 2c-f**, ORF1) but the shorter forms of ORF1 were not. The anti-X monoclonal antibody clearly detects 3 forms of ORF1 (**Fig 2d**, blue, green and orange arrowheads) whereas the polyclonal anti-X-Hel, RdRp or MetY antibodies revealed additional non-specific bands (**Fig 2 c, e, f**, asterisks), thus, rendering the identification of ORF1 shorter forms more difficult. Interestingly, and as previously observed [27], the anti-MetY WB showed an additional specific product migrating below 40 kDa, which could correspond to the predicted molecular weight of the Met subdomain (35 kDa, **Fig 2f**).

Together these results suggest that the ORF1 polyprotein is likely partially processed by cellular or viral proteases. However, the comparison of protein recognition profiles between wildtype (ORF1-WT and ORF1-V5-WT) and mutated ORF1 proteins (L1, L2, and L3) did not show any significant differences, indicating that the Hel/RdRp linker region does not modulate ORF1 protein maturation.

Next, to strengthen our observations and detect potential additional ORF1 cleavage products, an HA epitope was inserted downstream of the RdRp domain in ORF1-V5-WT and ORF1-V5-L1/L2/L3 constructs (**Fig 3a**, red arrowhead). At 24 h post-transfection, protein expression was analyzed by WB. As in **Fig 2**, WB with anti-V5, anti-X-Hel, anti-X, anti-RdRp, anti-MetY, and anti-HA antibodies showed similar recognition patterns between V5-HA-tagged wildtype and mutated ORF1 proteins (**Fig 3b-g**). The anti-HA antibody revealed two specific bands: one migrating at a size corresponding to the full-length ORF1, and another smaller band of lower intensity (**Fig 3g**, orange arrowhead). It should be noted that if cleavage were to occur between Hel and RdRp domains, a 51 kDa product corresponding to the predicted molecular weight of RdRp domain would be produced and specifically detected by anti-HA and anti-RdRp antibodies. The absence of detection of such a 51 kDa product in wildtype and mutant ORF1 proteins indicates that cleavage between Hel and RdRp domains is unlikely to occur.

**Fig 3.**
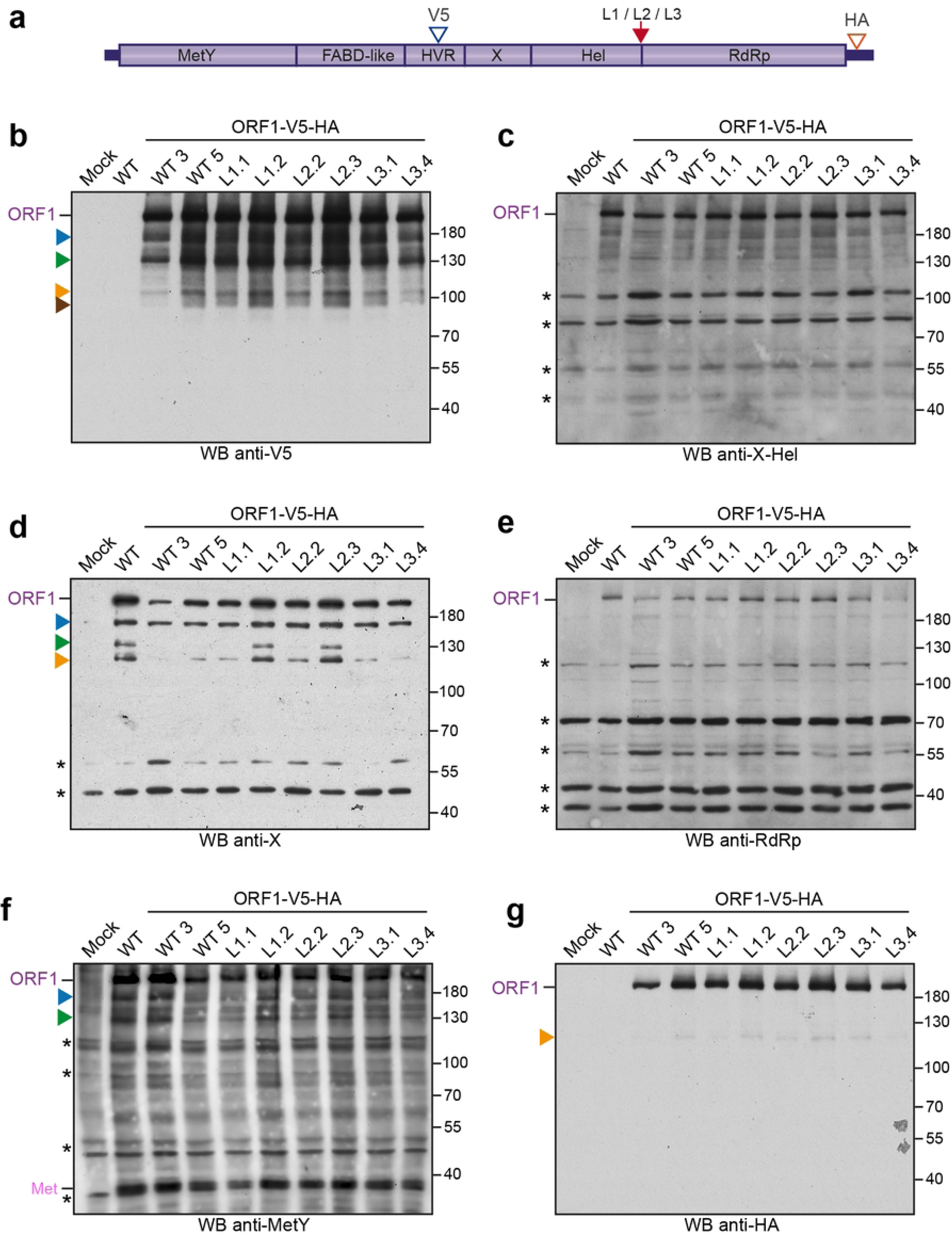
Western blot analysis of H7-T7-IZ cells expressing V5-HA-tagged wildtype and mutant ORF1 proteins in the Hel/RdRp linker region. (a) H7-T7-IZ cells were transfected with a pTM plasmid expressing the untagged (WT) or V5-HA-tagged wildtype ORF1 protein (V5 blue arrowhead, HA red arrowhead) or the ORF1 harboring mutations in the Hel/RdRp linker region (L1, L2, L3, red arrow). Total cell lysates were collected in the presence of protease inhibitors at 24 h post-transfection and analyzed by WB. Mock-transfected cells served as a negative control (Mock). Immunoblots were probed either with antibodies directed against the V5 epitope (b), the X-Hel domain (c), the macrodomain X (d), the RdRp domain (e), the MetY domain (f) or the HA epitope (g). The full-length ORF1 protein (ORF1) and the Met domain (Met) are indicated on the left. Proteins detected at lower molecular weights are also indicated (blue, green, orange and brown arrowheads). Molecular weight markers are indicated in kilodaltons on the right. Two clones have been analyzed per construct except for the untagged wildtype ORF1 (WT). Asterisks indicate non-specific bands.

Next, V5-HA-tagged wildtype and mutant ORF1 proteins were immunoprecipitated (IP) at 24 h post-transfection using anti-V5 or anti-HA antibodies to enrich for potential differentially processed products (**Fig 4**). No differentially processed ORF1 products were observed in either IP V5 or IP HA of cells expressing wildtype or mutated ORF1 proteins, further supporting the unlikeliness of a cleavage occurring within the Hel/RdRp linker region (**Fig 4e-h**). Interestingly, previously observed smaller forms of ORF1 were detected by the anti-HA and anti-V5 antibodies in IP V5 and IP HA (blue, green and orange arrowheads, **Fig 4e-h**), indicating that these products bear both epitopes. Moreover, an additional 60kDa product was detected by the anti-V5 antibody in IP V5 (**Fig 4e**, grey arrowhead). This band may correspond to the ORF1 protein missing both the MetY and Hel-RdRp domains.

**Fig 4.**
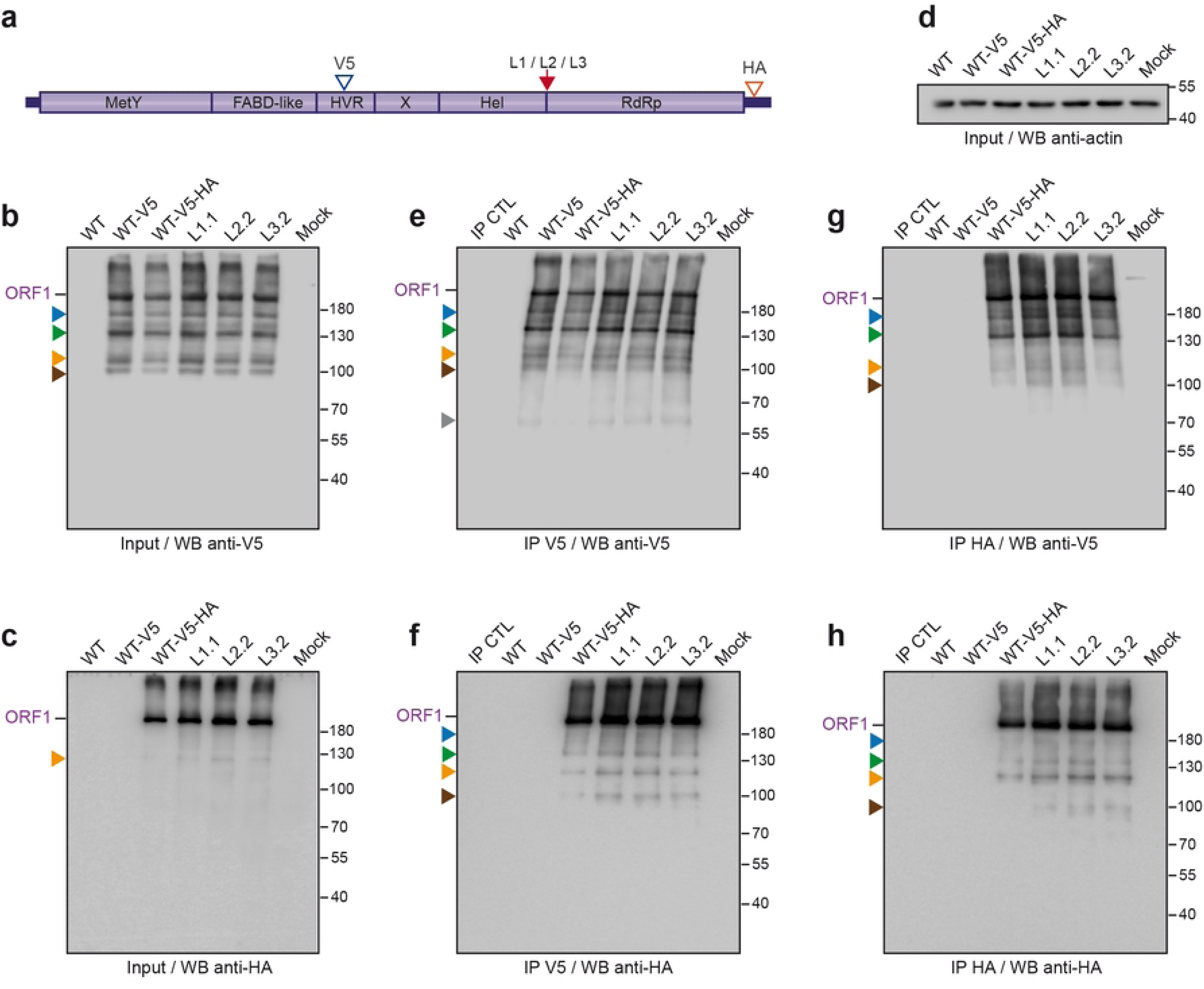
Immunoprecipitation of V5-HA-tagged wildtype and mutant ORF1 proteins in the Hel/RdRp linker region. (a) H7-T7-IZ cells were transfected with a pTM plasmid expressing the untagged (WT) or V5-HA-tagged wildtype ORF1 protein (V5 blue arrowhead, HA red arrowhead) or the ORF1 harboring mutations in the Hel/RdRp linker region (L1, L2, L3, red arrow). Total cell lysates were collected in the presence of protease inhibitors at 24 h post-transfection and immunoprecipitated (IP) using either anti-V5 (e, f) or anti-HA (g, h) antibodies. The expression levels of the V5-HA-tagged wildtype and mutant ORF1 proteins were checked in the inputs before IP by WB anti-V5 (b) and anti-HA (c). The anti-actin antibody was used to control for even loading (d). The full-length ORF1 protein (ORF1) is indicated on the right. Proteins detected at lower molecular weights are also indicated (blue, green, orange, brown and grey arrowheads). The wildtype or mutant ORF1 proteins were detected either with the anti-V5 (b, e, g) or the anti-HA (c, f, h) antibody. Magnetic beads alone (IP CTL) and mock-transfected cells (Mock) served as negative controls. Molecular weight markers are indicated in kilodaltons on the right.

Altogether, the absence of any minor cleavage product ascribable to either cleaved RdRp or Hel-containing but not RdRp-containing product and the lack of any effect from mutations in the Hel/RdRp linker region indicate that this disordered domain is not accessible to proteases for cleavage.

A recent study has shown that the ORF1 polyprotein interacts with the molecular chaperone HSP90 [28]. Inhibition of HSP90 by compounds such as NVP-HSP990 triggers ubiquitin-proteasomal degradation of ORF1, leading to the generation of a ∼120kDa C-terminally truncated ORF1 product, termed HEV-derived SMAD activator (HDSA), which promotes fibrogenic TGF-β signalling [29]. Accordingly, treatment of H7-T7-IZ cells expressing WT-V5-HA with MG132, NVP-HSP990, or both, revealed differential detection patterns of the full-length and ∼130kDa (**Figs 2-4**, green arrowhead) ORF1 products using anti-V5 antibody, suggesting that the latter corresponds to HDSA (**Fig 5b**, left). Detection with anti-HA (**Fig 5b**, right), anti-MetY (**Fig 5c**, top) and anti-X (**Fig 5c**, middle) antibodies confirmed degradation of the full-length ORF1 upon NVP-HSP990 treatment, as well as reduced HDSA accumulation upon MG132 treatment, as reported previously [28,29]. However, in our hands, NVP-HSP990 treatment did not lead to a detectable increase in HDSA accumulation compared with untreated cells. Interestingly, HDSA was differentially detected by the anti-MetY antibody (**Fig 5c**, top), confirming that this product is not processed at its N-terminus. In addition, Met subdomain detection was only slightly reduced upon NVP-HSP990 treatment, but strongly diminished by MG132 (**Fig 5c**, top), indicating that Met subdomain processing likely occurs independently of the ORF1-HSP90 complex.

**Fig 5.**
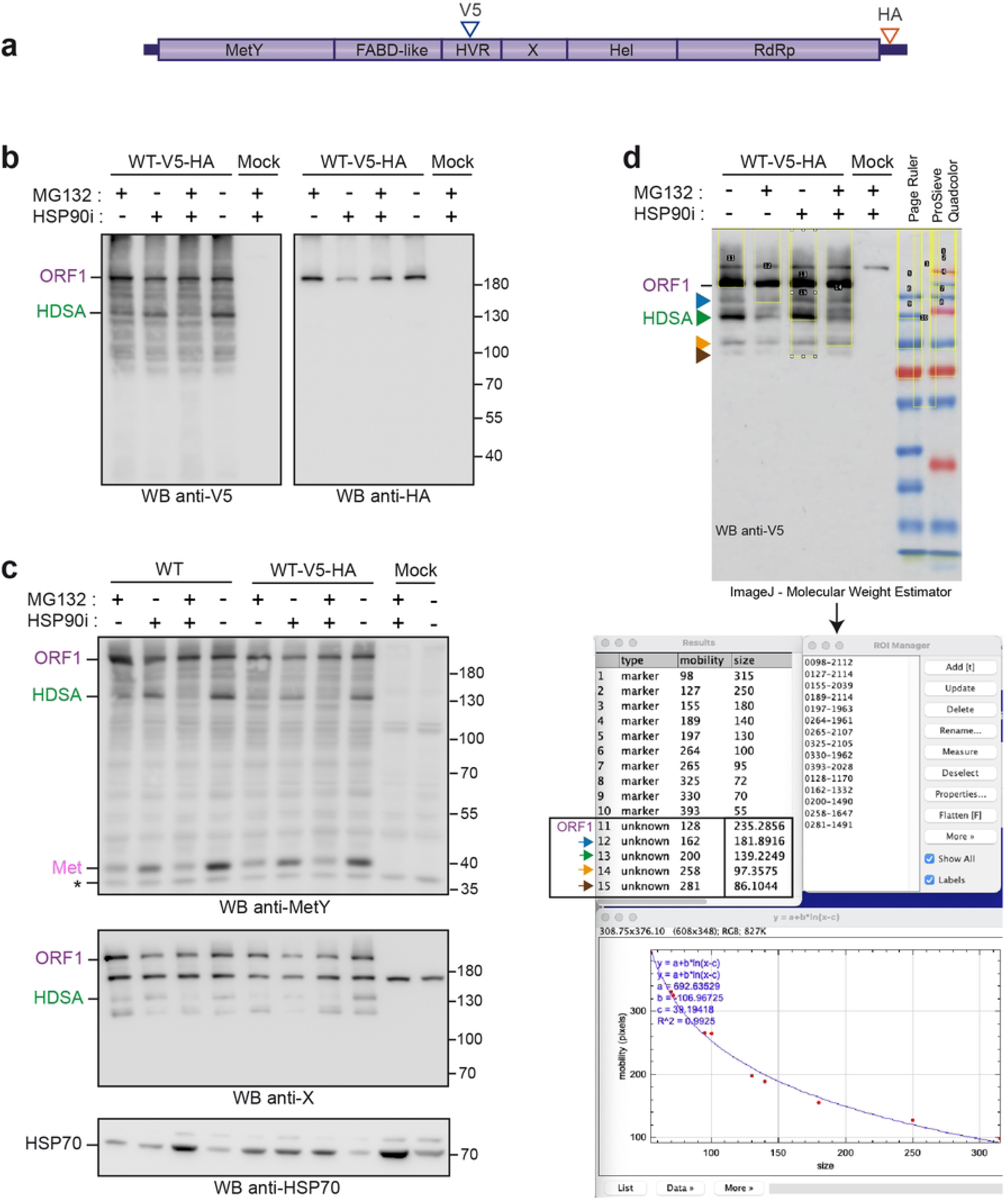
Effect of pharmacological inhibitors on ORF1 processing, and determination of the molecular weights of ORF1-derived products. (a) H7-T7-IZ cells were transfected with a pTM plasmid expressing the untagged (WT) or V5-HA-tagged wildtype ORF1 protein (V5 blue arrowhead, HA red arrowhead). (b, c, d) At 18h post-transfection, cells were treated with the proteasome inhibitor MG132 (10μM, +/-), the HSP90 inhibitor NVP-HSP990 (1μM, -/+), or both (+/+) for 9h. Cells treated with solvent (DMSO, -/-) and non-transfected cells (Mock) were used as controls. Total cell lysates were next collected in the presence of protease inhibitors and analyzed by WB. Immunoblots were probed either with antibodies directed against the V5 epitope (b, left), the HA epitope (b, right), the MetY domain (c, top), the macrodomain X (c, middle), or the HSP70 (c, bottom). The latter served as control for HSP70 expression, which is upregulated in response to chemical stress. The full-length ORF1 protein (ORF1), the HEV-derived SMAD activator (HDSA) and the Met domain (Met) are indicated on the left. Molecular weight markers are indicated in kilodaltons on the right. The asterisk on the WB anti-MetY indicates a non-specific band. (d) The *MolecularWeightEstimator* ImageJ macro [30] was used to determine the molecular weights of ORF1-derived products. A representative picture of WB anti-V5 is shown along with the colored protein markers used (Page Ruler and ProSieve Quadcolor). Regions of interest (ROI) were selected for both the marker bands of known size and the ORF1-derived bands of unknown size (ORF1 and blue, green, orange and brown arrowheads). The macro indexes their positions and calculates molecular weights based on a calibration curve fitted to the marker bands. Calculated sizes of full-length ORF1 and derived products are boxed. This analysis estimated the full-length ORF1 at ∼235 kDa, the blue arrowhead at ∼182 kDa, the green arrowhead/HDSA at ∼139 kDa, the orange arrowhead at ∼97 kDa, and the brown arrowhead at ∼86 kDa.

Finally, we applied an ImageJ macro (*MolecularWeightEstimator,* [30]) to more precisely determine the molecular weights of ORF1-derived products consistently observed in our anti-V5 WB (blue, green, orange and brown arrowheads in **Figs 2-4**). After selecting the known size markers and the unknown bands, the macro indexed their positions and computed their sizes from a calibration curve fitted to the markers (**Fig 5d**). Using this analysis, we estimated the full-length ORF1 at ∼235 kDa, the blue arrowhead at ∼182 kDa, the green arrowhead/HDSA at ∼139 kDa, the orange arrowhead at ∼97 kDa, and the brown arrowhead at ∼86 kDa.

#### Impact of mutations in the Hel/RdRp linker region on the ORF1 subcellular distribution

We next investigated the importance of the Hel-RdRp disordered region for the subcellular distribution of ORF1. To this end, cells expressing tagged wildtype and mutant ORF1 proteins were analyzed by immunofluorescence (IF) and confocal microscopy, at 24 h post-transfection. Anti-V5 (**Fig 6**, left panel) and anti-X-Hel (**Fig 6**, right panel) antibodies were used for V5-tagged constructs. Anti-V5 (**Fig 7**, left panel) and anti-HA (**Fig 7**, right panel) antibodies were used for V5-HA-tagged constructs. When ectopically expressed in H7-T7-IZ cells, the ORF1 protein displays both a diffuse cytoplasmic and dot-like distribution [23,31]. In **Fig 6** and **Fig 7**, wildtype (ORF1-V5-WT and ORF1-V5-HA-WT) and mutated ORF1 proteins (L1, L2, and L3) showed similar subcellular distribution profiles, regardless of the antibody used. Of note, HA staining revealed a similar pattern to that of V5 staining, indicating that the C-terminal epitope remains intact.

**Fig 6.**
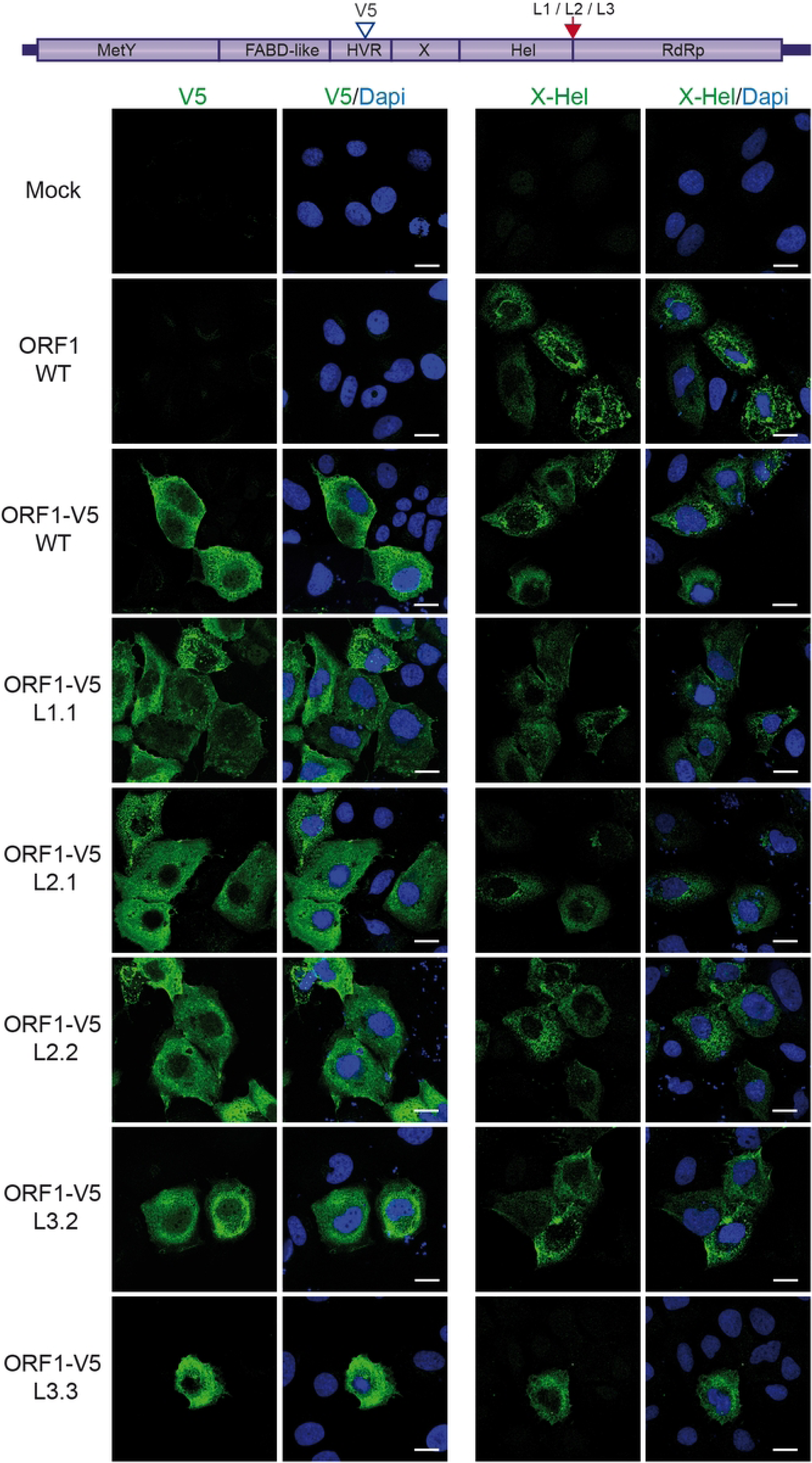
Subcellular distribution of V5-tagged wildtype and mutant ORF1 proteins in the Hel/RdRp linker region. H7-T7-IZ cells grown on coverslips were transfected with a pTM plasmid expressing the untagged (WT) or V5-tagged wildtype ORF1 protein (V5 blue arrowhead) or the ORF1 harboring mutations in the Hel/RdRp linker region (L1, L2, L3, red arrow). Two clones have been analyzed per mutant construct, except for L1 ORF1 mutant. Mock-transfected cells served as a negative control (Mock). Cells were fixed, permeabilized and stained 24h post-transfection. ORF1 proteins were stained using antibodies against V5 epitope (left panel) or X-Hel domain (right panel). Imaging was performed with a Zeiss LSM 800 confocal microscope, and fluorescence data were analyzed using ImageJ software.

**Fig 7.**
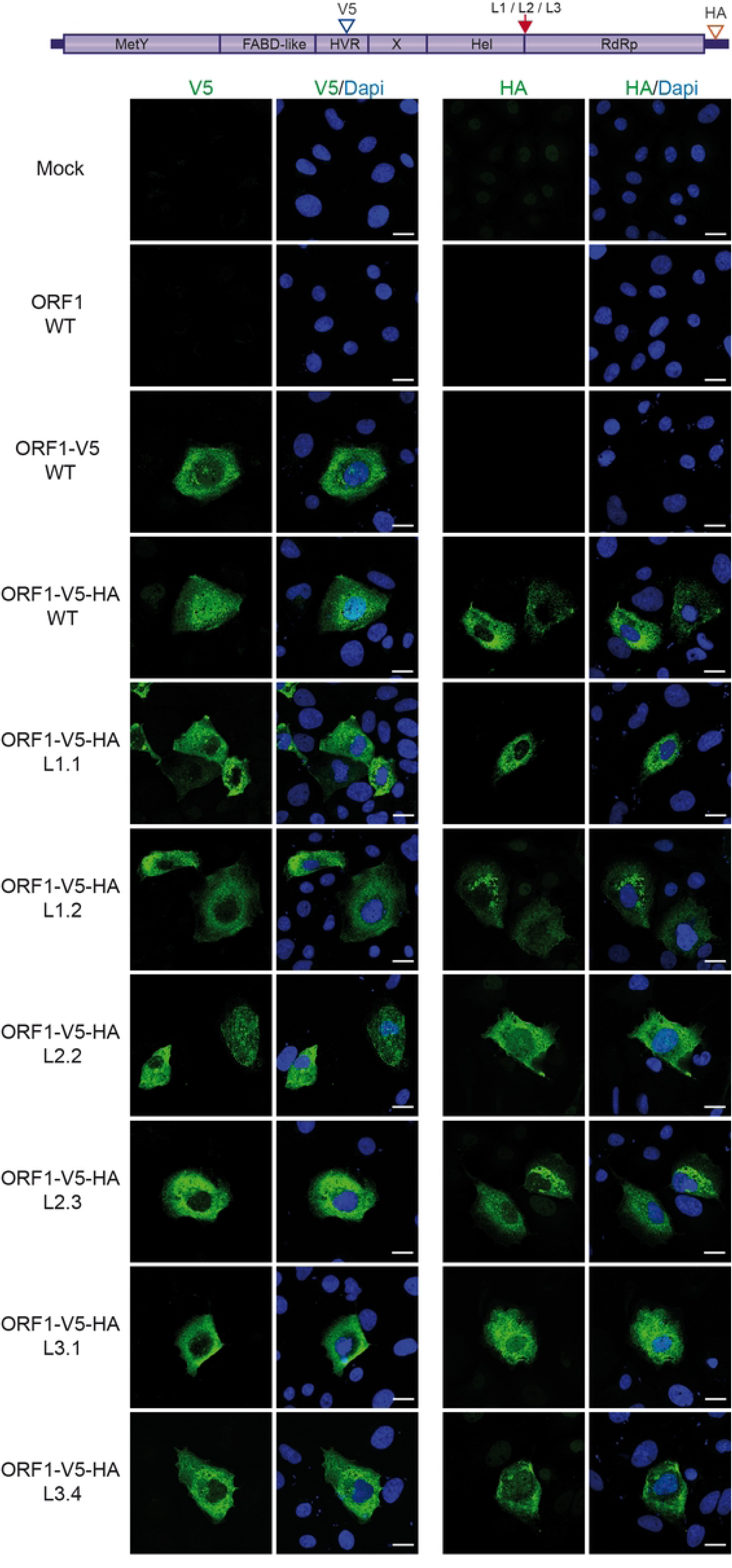
Subcellular distribution of V5-HA-tagged wildtype and mutant ORF1 proteins in the Hel/RdRp linker region. H7-T7-IZ cells grown on coverslips were transfected with a pTM plasmid expressing the untagged (WT) or V5-HA-tagged wildtype ORF1 protein (V5 blue arrowhead, HA red arrowhead) or the ORF1 harboring mutations in the Hel/RdRp linker region (L1, L2, L3, red arrow). Two clones have been analyzed per mutant construct. Mock-transfected cells served as a negative control (Mock). Cells were fixed, permeabilized and stained 24h post-transfection. ORF1 proteins were stained using antibodies against V5 (left panel) or HA epitope (right panel). Imaging was performed with a Zeiss LSM 800 confocal microscope, and fluorescence data were analyzed using ImageJ software.

These findings suggest that the mutations introduced into the Hel/RdRp linker region have no impact on ORF1 localization, indicating a lack of regulation in this region.

#### Impact of mutations in the Hel/RdRp linker region on viral replication

We next wondered whether mutations in the Hel/RdRp linker region could modulate HEV replication. For this purpose, we introduced the L1/L2/L3 mutations into an HEV replicon in which the ORF3 coding sequence and a part of ORF2 coding sequence were replaced by a *Gaussia* luciferase (GLuc) reporter gene. This tool enables replication efficiency to be measured by quantifying GLuc activity, which is dependent on ORF1-mediated transcription. A replication-deficient replicon (GLuc-ORF1-GAD), with a mutated GDD motif in the RdRp active site, was used as a negative control [25]. The luciferase activities were measured at 8, 24, 72, 96, 120 and 144 hours post-electroporation (p.e.) in the culture supernatant of PLC3 cells. For each time point, values are presented as fold increase compared to luciferase activities measured at 8 hours p.e. (**Fig 8a-c**).

**Fig 8.**
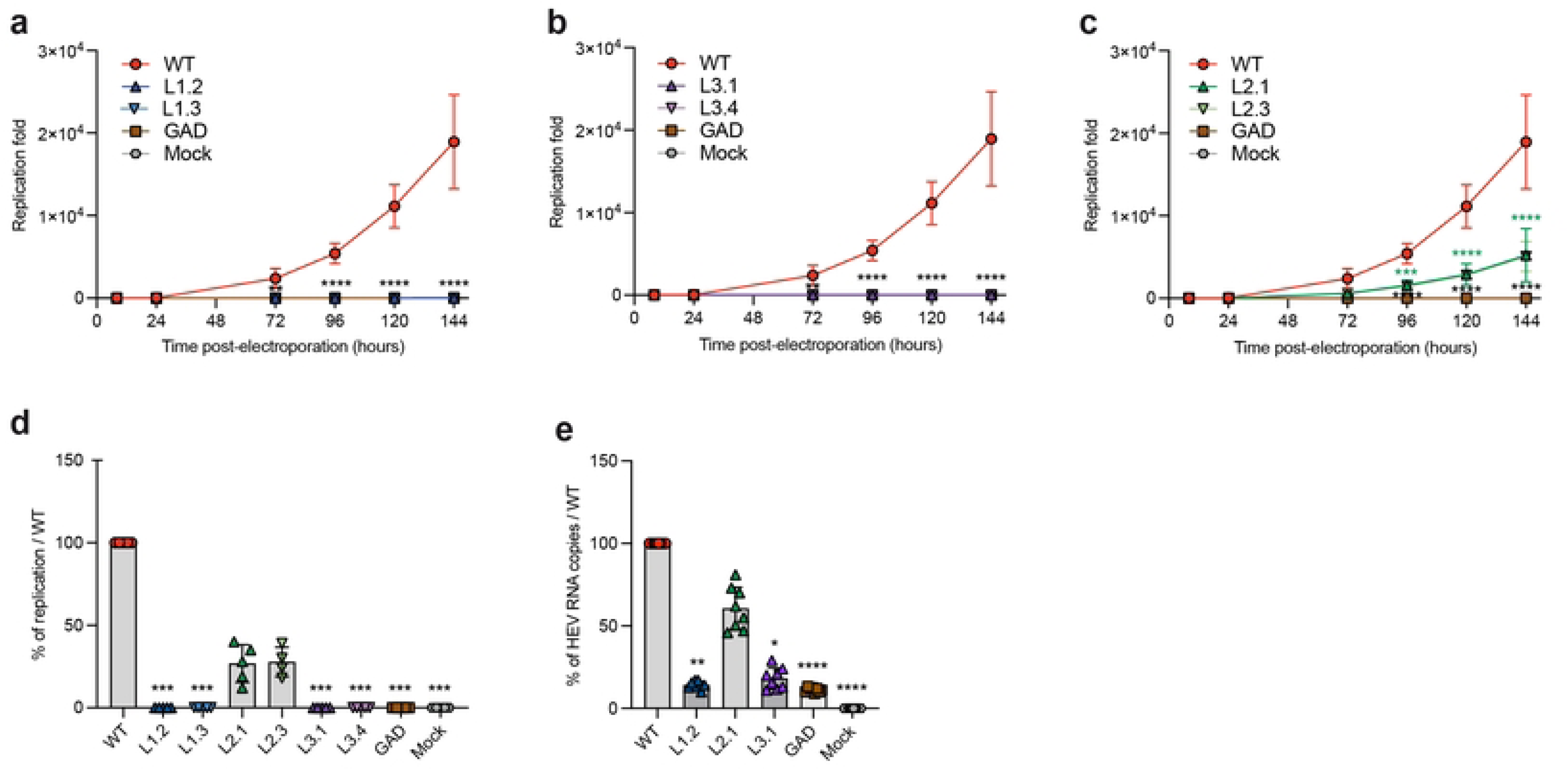
Replication efficacies of HEV-GLuc replicons expressing wildtype or linker-mutated ORF1. PLC3 cells were electroporated with HEV-p6-Gluc replicon constructs encoding the wildtype ORF1 protein (WT) or ORF1 mutated in the Hel/RdRp linker region. Supernatants were collected at 8, 24, 72, 96 and 144 hours post-electroporation (p.e.). Mock-electroporated cells (Mock) and a replication-deficient replicon (GAD) served as negative controls. Luciferase activity was measured by luminometry, normalized to the activity measured at 8 h p.e., and expressed as fold replication over time: mutant L1 (a), mutant L3 (b), and mutant L2 (c). A two-way ANOVA followed by Dunnett’s multiple comparison test was applied to assess differences between each mutant (L1, L2, L3) and WT (****p < 0.0001, 5 experiments, 3 repeats per experiment, 2 clones were analyzed per mutant, panels a–c). Results at 6 days (144 h) p.e. were also expressed as a percentage of WT replication efficacy (d), with significance tested using the Kruskal-Wallis test (***p < 0.001). For each mutant, numbers of genomic RNA copies were quantified by RT-qPCR using an ORF1 probe at 6 days p.e. (e) and results were expressed as percentages relative to the p6-GLuc WT replicon (****p < 0.0001, 4 experiments, 2 repeats per experiment, Kruskal-Wallis test).

The level of Gluc-ORF1-WT replicon increased steadily over time, reaching a 2 x 10^4^ fold increase at 144 hours p.e.. In contrast, the replication-deficient Gluc-ORF1-GAD replicon showed no replication activity. Interestingly, significant differences in replication efficiency between replicons carrying mutations in the Hel/RdRp linker region were observed. The L1 and L3 mutations resulted in a more than 95% reduction in replication efficiency at 144 hours p.e., compared with the Gluc-ORF1-WT replicon (**Fig 8d**). In contrast, replicons carrying L2 mutations increased steadily with time, reaching a 5.1 x 10^3^ fold increase at 144 hours p.e. (**Fig 8b**), corresponding to a 30% replication efficiency at 144 hours p.e. in comparison with the Gluc-ORF1-WT replicon (**Fig 8d**). We also quantified viral genomic RNA by RT-qPCR (**Fig 8e**). Consistently, the L2 mutant replicated with 60% efficiency at 144 hours p.e. compared to the Gluc-ORF1-WT replicon. Meanwhile, the L1 and L3 mutants exhibited replication efficiencies of 14% and 18%, respectively.

Taken together, our results indicate that the linker region between the Hel and RdRp domains of ORF1 is probably not involved in the regulation of ORF1 maturation and subcellular distribution, but is likely instrumental for HEV genome replication.

### Functional analysis of the C-terminal tail of the RdRp domain on viral replication

We next investigated whether other mechanisms could regulate the HEV viral replicase. Based on structural modelling with AlphaFold2, we proposed that the ORF1 protein may self-regulate through conformational changes in different regions [11]. Specifically, the 20 C-terminal residues of the RdRp domain (residues 1746-1765 in HEV-p6) appeared capable of adopting two distinct conformations, giving rise to alternate folds of the ‘fingertips’, the structural elements connecting the fingers and thumb domains in viral RdRps (**Fig 1a**, bottom). The structural divergence originates at residue F1746, which anchors the C-terminus within a hydrophobic pocket of the thumb. In one conformation, the F1746 residue is fully inserted into this pocket, exposing downstream residues, including the C-terminal helix 1757-1765, at the fingertips’ surface (**Fig 1a**, bottom left). In the alternate conformation, the interaction between the F1746 residue and the thumb’s pocket is weakened, and residues 1747-1752 instead form a beta-strand within the fingertips, thereby repositioning helix 1757-1765 to the opposite side of the fingertips (**Fig 1a**, bottom right). This conformational plasticity within a region critical for RdRp activity [32] suggests that the activity of HEV RdRp may be subjected to regulation by its C-terminal region.

To test this hypothesis, we generated the ORF1 ΔC-ter1 construct in which the 19 last C-terminal residues of RdRp were deleted by inserting a stop codon downstream of F1746 (**Fig 1b**, bottom). The impact of this deletion on HEV replication was assessed using an HEV-p6-GLuc replicon during a 144 hours kinetics, as described above. The ΔC-ter1 replicon showed a drastic reduction in replication efficiency in PLC3 cells as measured by luminometry (**Fig 9a**). At 144 hours p.e., replication was completely abolished in comparison with the wildtype replicon (**Fig 9b**), a finding that was corroborated by quantifying replicon genome copy numbers by RT-qPCR (**Fig 9c**). Contrary to our initial hypothesis, deleting the C-terminal region of RdRp did not result in a constitutively more active polymerase; rather, it was detrimental to HEV replication in PLC3 cells. These results suggest that the RdRp C-terminus is essential for HEV replication.

**Fig 9.**
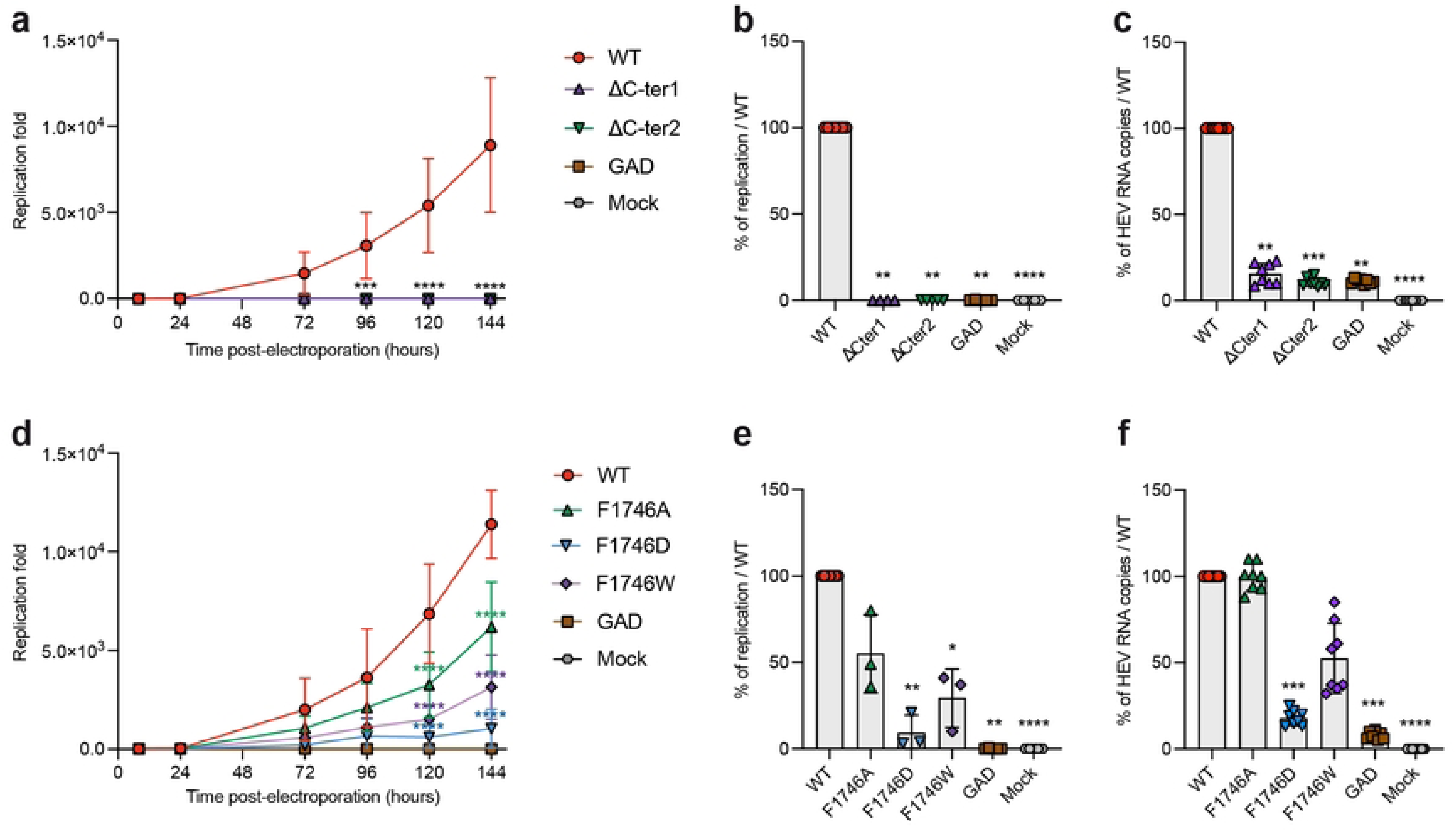
Replication efficacies of HEV-GLuc replicons expressing wildtype or C-terminus-mutated ORF1. PLC3 cells were electroporated with a Gluc replicon system encoding for the wildtype ORF1 protein (WT) or ORF1 mutated in RdRp C-ter. Supernatants were collected at 8, 24, 72, 96, and 144 hours post-electroporation (p.e.). Mock-electroporated cells (Mock) and a replication-deficient replicon (GAD) served as negative controls. Luciferase activity was measured by luminometry, normalized to the activity measured at 8 h p.e., and expressed as fold replication over time: mutants ΔC-ter1 and ΔC-ter2 (a), soft mutations F1746A/D/W (d). A two-way ANOVA followed by Dunnett’s multiple comparison test was applied to assess differences between each mutant and WT (*****p < 0.0001, 3 experiments, 3 repeats per experiment, panels a and d). Results at 6 days (144 h) p.e. were also expressed as a percentage of WT replication efficacy for ΔC-ter mutants (b) and soft mutants F1746A/D/W (e), with significance tested using the Kruskal-Wallis test (***p < 0.001, panels b and e). For each ΔC-ter (c) and soft mutant (f), numbers of genomic RNA copies were measured by RT-qPCR using an ORF1 probe at 6 days p.e. and results were expressed as percentages relative to the p6-GLuc WT replicon (****p < 0.0001, 3 experiments, 4 repeats per experiment, Kruskal-Wallis test).

We next tried milder modifications than the introduction of a stop codon downstream of the F1746 residue (**Fig 1b**, bottom). First, we generated the ORF1 ΔC-ter2 construct in which a stop codon was introduced at position 1757, thus deleting a shorter fragment consisting of the last 8 residues that constitute the terminal α-helix. This mutation enabled us to examine the impact on replication without modifying the region interacting with the RdRp hydrophobic pocket. Moreover, we examined the impact of altering the balance between the two putative conformations on replication. We introduced several mutations to destabilize the interaction between the C-terminal and N-terminal regions of RdRp to varying degrees. The F1746A mutation was expected to moderately destabilize the C-terminal region, favouring the weakest interaction. In contrast, the F1746D mutation was predicted to severely destabilize the C-terminal region of RdRp, thereby preventing interaction with the RdRp N-terminal region. Lastly, the F1746W mutation was expected to strengthen the interaction between the C-terminal region and the rest of ORF1. None of the mutants enhanced HEV replication, but replication was reduced to varying extents by the different mutations: the ΔC-ter2 mutation was as deleterious to HEV replication as the ΔC-ter1 (**Fig 9a-c**), suggesting a direct role of the C-terminal α-helix at the replication step. The point mutants of F1746 were milder, with F1746A, F1746D and F1746W reducing replication by approximately 50%, 90% and 75%, respectively, in comparison with the wildtype replicon (**Fig 9d-e**). Interestingly, the effects of mutations on viral replication, as measured by RT-qPCR, contrasted with those measured by luminometry. Indeed, the F1746A mutation did not affect the number of genomic viral RNA copies, while the F1746D and F1746W mutations decreased genome replication (**Fig 9f**), but not as much as measured by luminometry (**Fig 9e**). This suggests that the F1746 residue may play a specific role in transcription of HEV subgenomic RNAs as well as in replication.

Together, our results highlight a critical role of the 20 C-terminal amino acid residues in RdRp activity. This region appears to be intolerant of sequence modifications, since both complete (ΔC-ter1) and partial (ΔC-ter2) deletions of the C-terminal region resulted in the loss of polymerase activity. Furthermore, mutations at position F1746 (except F1746A on genome copy numbers) resulted in reduced replication, suggesting that regulation of the contact between the C-terminus and the thumb’s hydrophobic pocket of RdRp is likely essential for viral replication.

## Discussion

Although replication is an essential step in the HEV lifecycle, the processing and structural organization of the HEV replicase remain controversial. All studies have shown that ORF1 was mainly expressed as a full-length protein, but some authors have also reported products of smaller size that may represent processed fragments of ORF1 (reviewed in [19]). The processing of non-structural polyproteins is often a key process in the lifecycle of viruses since it leads to the production of functional proteins that regulate the viral genome synthesis. Still, in animal positive-sense RNA viruses, there are known cases of non-processed replication polyprotein, such as for the Flock House Virus (FHV, [33]). Like the FHV protein A and many plant replicases [34,35], HEV ORF1 does not encode a protease [11,15]. This could explain why most ORF1 proteins remain unprocessed, and the discrepancies in reports on ORF1 partial processing as it may depend on a cellular protease not equally available in the various expression systems used. We have previously shown that ORF1 is expressed as a full-length protein, but also as smaller proteins of lower intensity (95-170 kDa) using three different systems expressing the full-length protein [23]. In addition, analysis by nano-scale liquid chromatography coupled to tandem mass spectrometry suggested that the N-terminus of ORF1 is likely conserved in smaller size ORF1 proteins [23]. This observation is in accordance with recent studies reporting that the ubiquitin-proteasomal degradation of ORF1 leads to the generation of HDSA, a ∼120 kDa C-terminally truncated ORF1 product [28,29].

In the present study, we characterized V5-or V5-HA-tagged-ORF1 proteins expressed in Huh-7 cells. Using an extensive antibody panel targeting the V5 tag, HA tag, X-Hel domain, X domain, RdRp domain and MetY domain, we confirmed that ORF1 is predominantly expressed as a full-length protein, together with several truncated products of lower abundance. Notably, WB using the monoclonal anti-V5 antibody, combined with a precise molecular weight determination tool, revealed the full-length ORF1 protein migrates at an apparent molecular weight of ∼235 kDa, as well as ORF1-derived products of apparent molecular weights of ∼182 kDa (blue), ∼139 kDa (green), ∼97 kDa (orange) and ∼86 kDa (brown) (**Figs 2b, 3b, 4b** and **5d**). Of note, the calculated molecular weight of full-length ORF1 markedly differs from its theorical value (193.7 kDa), suggesting that (i) ORF1 undergoes post-translational modifications such as polyubiquitination, (ii) ORF1 exhibits aberrant migration in SDS-PAGE due to its membranous regions, or (iii) both mechanisms contribute. In addition, the use of pharmacological inhibitors revealed that the ∼139 kDa ORF1 product (green) corresponds to the recently reported HDSA [28,29] (**Fig 5b-c**). The same recognition pattern, except for the ∼86 kDa product (brown), was also detected by the anti-X monoclonal antibody (**Figs 2d** and **3d**). The V5 epitope was inserted in the HVR, just upstream of the macrodomain X. Accordingly, antibody binding sites may be located in similarly exposed regions of ORF1, providing access to both V5 and X epitopes and explaining the overlapping recognition profiles. In contrast, the full-length ORF1 protein was the predominant species detected with the anti-X-Hel (**Figs 2c** and **3c**) and anti-RdRp (**Figs 2e** and **3e**) polyclonal antibodies. The ∼182 kDa product (blue) was also recognized by the anti-X-Hel antibody in V5-tagged ORF1 expressing cells (**Fig 2c**), but was barely detectable in cells expressing HA-V5-tagged ORF1 (**Fig 3c**). The low levels of truncated ORF1 products, combined with the lower signal-to-noise ratio of the anti-X-Hel and anti-RdRp polyclonal antibodies, may account for the difficulty in consistently detecting these products. Finally, the monoclonal anti-HA antibody clearly detected the full-length ORF1 protein, as well as a much weaker ORF1 product likely corresponding to the ∼97 kDa ORF1-derived protein (orange) (**Fig 3g**). The affinity of the antibody as well as the localization of its epitope at the C-terminus of the ORF1 protein may explain this recognition pattern.

Despite the distinct reactivity and specificity observed for the 6 different antibodies used (**Figs 2** and **3**), no differences in recognition profiles or signal intensity were detected when comparing total protein extracts from cells expressing either wildtype ORF1 or mutants engineered in the Hel-RdRp linker region. To increase the likelihood of detecting low-abundance products, we exploited the double-tagged ORF1 construct for immunoprecipitation experiments (**Fig 4**). Regardless of the antibody used for IP or blot detection, protein profiles were identical between wildtype and all 3 linker mutants, suggesting that cleavage within the Hel-RdRp linker is unlikely. However, based on the predicted molecular weights of individual ORF1 domains, several hypotheses can be drawn from the IP results.

First, the ∼182 kDa ORF1 product (blue), detected by both anti-HA and anti-V5 antibodies in inputs and IP, may represent an ORF1 protein lacking the Met subdomain. Supporting this hypothesis, a ∼30 kDa product was consistently detected with the anti-MetY polyclonal antibody (**Figs 2f, 3f** and **5c**, Met). Moreover, AlphaFold2 structural modeling of this region revealed a small solvent-exposed segment comprising two successive helix-loop motifs of unknown function, positioned between the Met and Y subdomains (**S1a Fig**, red). This structurally distinct region, not clearly assignable to either Met or Y, may be particularly susceptible to proteolytic cleavage or degradation. Consistent with this hypothesis, detection of the Met subdomain was reduced in the presence of the proteasome inhibitor MG132 (**Fig 5c**), which at high concentrations also inhibits lysosomal cathepsins and cytosolic calpains [36–39]. This finding is in agreement with a previous study showing that Met domain processing depends on cysteine proteases [27].

Based on its molecular weight, proteasome dependency, and recognition by different antibodies (**Fig 5**), we hypothesized that the ∼139 kDa ORF1 product (green) corresponds to HDSA, a C-terminally truncated ORF1 product lacking the Hel and RdRp domains [28,29]. Unexpectedly, this product was detected by both anti-V5 and anti-HA antibodies in IP experiments (**Fig 4**). This observation raises the possibility that the ∼139 kDa band may represent a mixture of distinct products of similar size, warranting further investigation.

The detection of ∼97kDa (orange) and ∼86 kDa (brown) ORF1 products by both anti-HA and anti-V5 antibodies in all IP experiments (**Fig 4**) suggests the occurrence of additional cleavage events, one between the FABD-like domain and the HVR region, and another within the HVR region just upstream of the V5 epitope site (**S1b Fig**). Finally, the approximately 60kDa ORF1 product (grey), observed exclusively following IP V5 and detection with the anti-V5 antibody (**Fig 4e**), may represent the Met-truncated N-terminal fragment of ORF1 or HDSA. This interpretation is consistent with the distribution of unstructured residues within the ORF1 N-terminus of ORF1: the longest stretch corresponds to the HVR (>70 residues), followed by an exposed region between the Met and Y subdomains (49 residues) and a shorter linker connecting MetY to FABD (10 residues) [11].

Together, our results show that the ORF1 polyprotein is predominantly expressed as a full-length protein, together with several lower-abundance truncated products, including the previously described HDSA [28,29]. Mutations introduced within the Hel-RdRp linker did not alter ORF1 processing, indicating that cleavage in this region is unlikely. Beyond the proteasomal processing up to the X domain that generates HDSA [29], our data suggest that the ORF1 polyprotein may also undergo additional cleavages between the Met and Y regions, and within the C-terminal extension of the FABD domain or within the HVR (**S1b Fig**). However, we cannot exclude the possibility of further proteasomal processing or spontaneous truncations/breakages occurring in structurally fragile or exposed regions. These hypotheses will require further validation, notably by assessing the impact of mutations introduced in other linker regions on the expression of ORF1 products.

Regarding the subcellular distribution of the Hel-RdRp linker mutants, no significant differences were observed compared to the wildtype ORF1, as shown by labeling V5- and HA-V5-tagged ORF1 with three different antibodies targeting V5, HA and the X-helicase domain (**Figs 6** and **7**). In contrast, analysis using the GLuc HEV replicon system revealed a strong replication defect: replication was almost completely abolished in the L1 and L3 mutants, and significantly reduced in the L2 mutant (**Fig 8**). Conservation analysis of the linker region, based on an alignment of 50 reference HEV sequences [40], showed that three glycines out of six mutated residues in L1 were highly conserved (**S1c Fig)**. For L2 and L3 mutants, three of the five mutated residues were also highly conserved, corresponding to the RPS and PRG motifs, respectively (**S1c Fig**). These findings suggest that amino acid substitutions in this region may alter the conformation and/or relative flexibility of the Hel and RdRp domains, thereby impairing replication. The more pronounced defects observed in the L1 and L3 mutants compared with L2 may be explained by the location of the mutated residues near the C-terminus of the Hel domain (L1) and the N-terminus of the RdRp domain (L3), where they may directly affect domain function. In summary, the Hel-RdRp linker region of HEV ORF1 does not appear to harbor cleavage sites. However, the strong evolutionary conservation of residues within this disordered region underscores its essential role in maintaining efficient viral genome replication.

Positive-sense RNA viruses that express their replicase as a polyprotein face the general challenge of avoiding excess RdRp production. They regulate the number of active RdRp molecules by various means, *e.g.* degradation of most RdRp molecules after release from the polyprotein [41]. This may indeed be one function of proteasomal degradation of the C-terminus of the ORF1 polyprotein to generate HDSA [29] and shorter fragments. From our previous modelling of ORF1 and our present observation that the HEV RdRp is not detectably cleaved from ORF1, we hypothesize that two layers of regulation through flexibility may be at work in regulating the HEV RdRp activity: (i) flexibility between the Hel and RdRp domains (see above), and (ii) alternative folding of the RdRp C-terminus (**Fig 1a**, bottom).

The HEV RdRp appears to adopt an unusually complex fold compared with other positive-sense RNA virus RdRps. In canonical viral RdRps, the ‘fingertips’, the part of the enzyme connecting the N-terminal ‘fingers’ domain to the C-terminal ‘thumb’ domain, are formed solely by N-terminal segments of the RdRp (two of which are shown on **Fig 1a**, colored in blue and cyan, bottom). In contrast, in the HEV RdRp, the fingertips have an extra segment brought by the very C-terminus of HEV RdRp. Its alternate conformations may reflect difficulties for AlphaFold in completing such a complex fold, but may also reflect an intrinsic propensity of the HEV RdRp to misfold. If so, the ORF1 molecules where RdRp is misfolded may be the prime targets of the ubiquitin-proteasome system.

Functionally, our results demonstrate that the 20 C-terminal residues of ORF1/RdRp (in red on **Fig 1a**, bottom) are critical for RNA synthesis. Deletion of as few as the last eight residues completely abolishes replication in the GLuc replicon system. These findings suggest that the conformation in which these residues integrate into the fingertips (**Fig 1a**, bottom right) likely corresponds to the active RdRp fold. In contrast, the alternate conformation (**Fig 1a**, bottom left) is probably inactive. Our results also pinpoint the residue F1746 as a key regulator of RdRp activity. F1746 anchors the C-terminal segment in the RdRp’s thumb just upstream of its engagement in the fingertips. Replacing F1746 with a bulkier aromatic residue (F1746W), presumably strengthens the hydrophobic anchoring interaction, and is deleterious to replication. However, weakening the interaction (F1746A) or abrogating it (F1746D) is also deleterious. Interestingly, we also found that this interaction may be particularly important for the transcription and/or translation of the subgenomic RNAs (**Fig 9**), which warrants further investigation. Taken together, our results suggest that the C-terminal segment and its anchor in the thumb are important regulatory elements of RdRp activity.

## Acknowledgments

We thank Suzanne U. Emerson (NIH, USA), Jérôme Gouttenoire (University of Lausanne, Switzerland), Ralph Bartenschlager (University of Heidelberg, Germany), Tero Ahola (University of Helsinki, Finland), and Qiang Ding (Tsinghua University, China) for providing us with reagents. We thank Jean Dubuisson (Center for Infection and Immunity of Lille, France) for critical reading of the manuscript. We would also like to thank the imaging core facility of the BioImaging Center Lille-Nord de France (US 41 - UAR 2014 - PLBS) for access to the infrastructure, and the integrative bioinformatics BIOI2 facility for making the ColabFold pipeline easily accessible at I2BC.

## Author Contributions

Material preparation, experiments, data collection and analysis were performed by L.M., S.F., C.M., M.F., T.T., G.V.A., S.B., L.C., and C-M.A-D.

Writing the original draft of the manuscript was performed by L.M., L.C. and C-M.A-D

Reviewing and editing the manuscript were performed by L.M., S.F., M.F., S.B., L.C. and C-M.A-D.

All authors read and approved the final manuscript.

---

**Sup 1 Fig.**
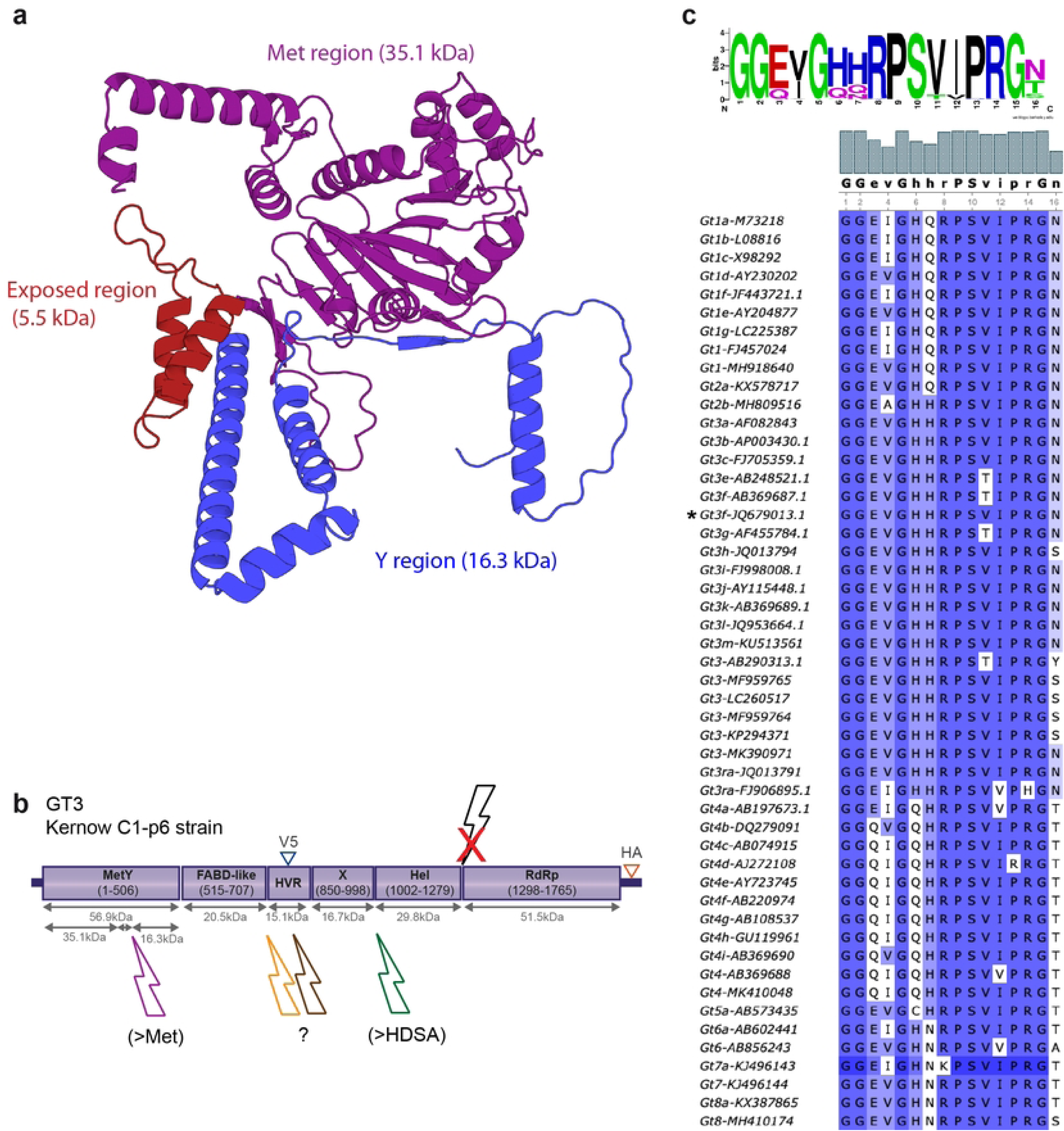

